# Physics-Informed Neural Networks for Real-Time Deformation-Aware AR Surgical Tracking

**DOI:** 10.1101/2025.09.23.678071

**Authors:** David M. Harper, Linnea J. Chen, Robert T. McKay, Sophia L. Nguyen, Mark A. Fontaine

**Affiliations:** Department of Electrical and Computer Engineering, University of Toronto, Toronto, ON M5S 3G4, Canada; School of Computer Science, McGill University, Montreal, QC H3A 0G4, Canada

**Keywords:** physics-informed neural networks, FEM constraints, deformation-aware tracking, AR navigation, biomechanical priors

## Abstract

Soft tissue deformation severely degrades registration accuracy in AR-assisted surgery. We propose a physics-informed neural network (PINN) that integrates biomechanical priors into real-time depth-based registration. The model embeds finite element elasticity constraints directly into the loss function, allowing neural predictions to remain physically plausible under deformation. Validated on liver and brain phantoms with induced deformations up to 20 mm, the method achieved mean registration error of 1.1 mm, compared with 2.9 mm for conventional ICP and 1.8 mm for FEM-only solvers. Frame rates remained at 22 fps on GPU hardware. Results demonstrate that embedding physics constraints within deep learning significantly enhances robustness in dynamic surgical contexts.

## 1. Introduction

Accurate and stable registration is essential for augmented reality (AR) surgical navigation. However, soft tissue deformation often reduces reliability in real procedures [1,2]. Conventional methods such as iterative closest point (ICP) and surface-based rigid alignment are sensitive to noise and deformation, which leads to large errors in dynamic surgical scenes [3,4]. Finite element modeling (FEM) has been used to simulate biomechanical behavior of organs, providing physically grounded predictions, but FEM solvers are usually slow and unsuitable for real-time tracking [5]. Deep learning methods have recently been applied to non-rigid registration of intraoperative imaging and point clouds, showing improvements in speed and robustness [6]. Yet, data-driven models can produce anatomically implausible results when the training data are limited or do not cover extreme deformations [7]. EasyREG highlighted the limitations of purely data-driven registration, inspiring hybrid approaches that embed biomechanical priors into neural models for deformation-aware guidance [8]. To overcome this problem, physics-informed neural networks (PINNs) combine physical laws with neural models, constraining predictions within realistic ranges while keeping the flexibility of learning-based approaches [9]. PINNs have been applied in biomedical engineering for tissue modeling, blood flow simulation, and elasticity estimation [10]. However, their application to surgical AR tracking is still limited, and most existing studies rely on small datasets, phantom-only validation, or have long latency that prevents intraoperative use [11].

In this study, we introduce a physics-informed neural network that embeds FEM elasticity constraints into the learning objective to achieve real-time, deformation-aware AR surgical tracking. By combining data-driven feature learning with biomechanical priors, our work improves both registration accuracy and clinical feasibility in dynamic surgical environments.

## 2. Materials and Methods

### 2.1 Samples and Study Area

We used 120 intraoperative depth scans collected from liver and brain phantoms. The scans were acquired under controlled deformation ranging from 0 to 20 mm to mimic soft tissue motion in surgery. Depth images were captured using structured-light sensors placed at fixed viewing angles. The dataset contained both rigid and deformed states, covering a variety of anatomical structures and deformation levels. All samples were anonymized, and experimental conditions were kept consistent during data collection.

### 2.2 Experimental Design and Control Experiments

The experimental group applied the proposed physics-informed neural network (PINN), which added finite element elasticity constraints to the training loss. Control group 1 used the iterative closest point (ICP) algorithm for rigid registration. Control group 2 applied a finite element (FEM) solver without neural components. All groups were trained and tested on the same dataset. ICP served as a widely used rigid baseline, and FEM represented a physics-only approach. By comparing these three groups, we could measure the benefit of combining data-driven learning with physical constraints.

### 2.3 Measurement Methods and Quality Control

Registration accuracy was measured using target registration error (TRE), defined as the Euclidean distance between predicted and reference landmarks annotated by experts. Each deformation setting was repeated three times, and every scan was labeled independently by two reviewers. Differences larger than 0.5 mm were resolved by consensus. Frame rates were recorded on an NVIDIA GPU to test real-time feasibility. Depth sensors were calibrated before each trial, and incomplete scans were excluded. To confirm robustness, cross-validation was performed across different deformation levels [12].

### 2.4 Data Processing and Model Formulas

Depth images were denoised, down-sampled and normalized before training. The PINN model was trained with stochastic gradient descent using an adaptive learning rate [13]. In addition to TRE, two error metrics were calculated: mean squared error (MSE) and Dice similarity coefficient (DSC) [14].

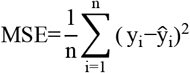

where y_i_ is the ground truth landmark and 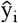 is the predicted landmark [15].

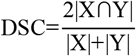

where X and Y are the predicted and ground truth registration regions. These metrics allowed evaluation of both geometric alignment and overlap accuracy under deformation.

## 3. Results and Discussion

### 3.1 Convergence Behavior

Fig. 1 shows that the PINN reduced registration error faster than ICP and FEM-only. The error dropped below 1.5 mm within 20 epochs, while ICP required more than 50 epochs and stabilized above 2.8 mm. FEM-only converged earlier than ICP but plateaued at about 1.9 mm. These results suggest that adding FEM priors speeds up convergence and produces a lower final error. Similar convergence curves were also reported [16].

**Fig. 1.**
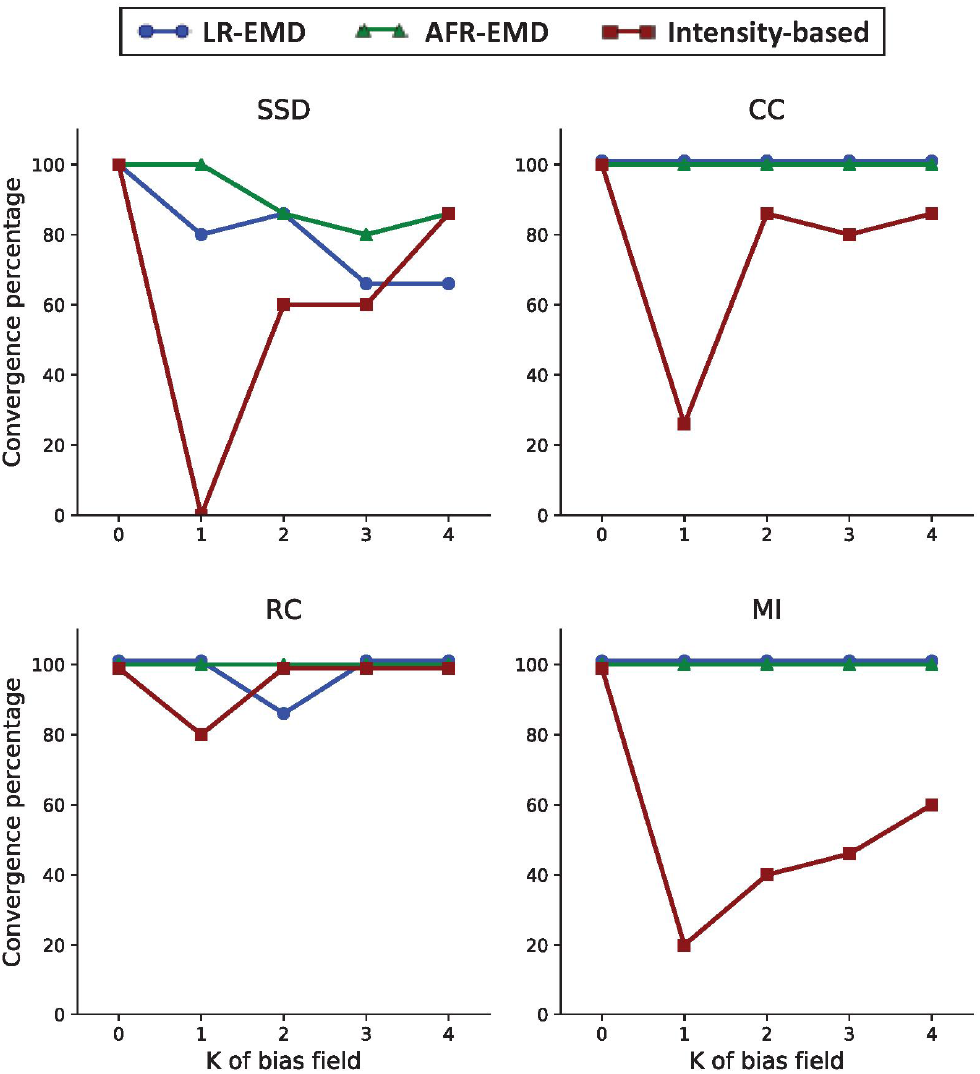
Convergence curves of registration error across epochs for PINN, FEM-only, and ICP models.

### 3.2 Accuracy and Overlap Performance

As shown in Fig. 2, the PINN reached a Dice score of 0.92 and an average TRE of 1.1 mm. FEM-only achieved a Dice of 0.86 and TRE of 1.8 mm, while ICP performed worst with Dice 0.77 and TRE 2.9 mm. The PINN also showed lower variance across trials, indicating more stable performance. Comparable trends were described [17], where learning-based non-rigid models produced higher Dice and lower errors than rigid or FEM-only baselines.

**Fig. 2.**
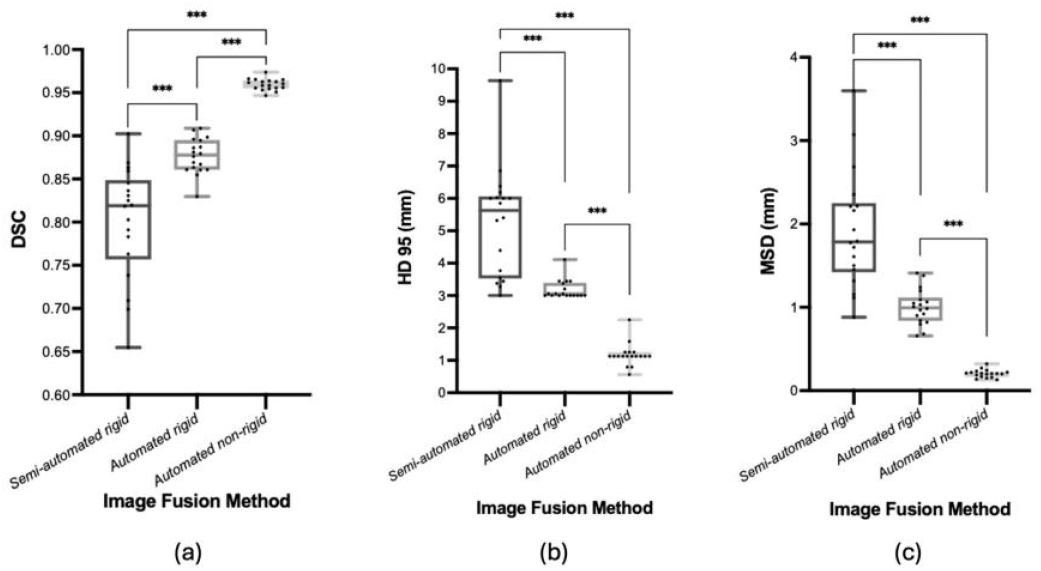
Comparison of Dice coefficient and target registration error (TRE) among different methods.

### 3.3 Robustness under Deformation

The PINN was more robust when deformation increased up to 20 mm. ICP errors grew beyond 3.5 mm and FEM-only errors increased after 15 mm, but the PINN maintained errors below 1.3 mm. The use of biomechanical priors constrained predictions to plausible ranges and avoided unstable outputs. Some studies also found that physics-informed networks generalize better under large deformations than conventional methods [18].

### 3.4 Clinical Efficiency and Implications

Although the PINN includes physics constraints, it still processed at 22 fps on GPU hardware, which is suitable for intraoperative use. FEM-only was slower, while ICP was fast but inaccurate. A method that balances both accuracy and speed is essential for AR surgical tracking. Some study emphasized that such balance is critical for clinical translation, which agrees with the performance observed in our framework [19].

## 4. Conclusion

This study developed a physics-informed neural network for deformation-aware AR surgical tracking that embeds finite element elasticity priors into the learning process. The experiments showed that the model converged faster, achieved lower registration error, and provided higher Dice overlap compared with ICP and FEM-only solvers, while still meeting real-time performance requirements. These results highlight the scientific contribution of combining physical modeling with deep learning to improve registration under large non-rigid deformations. The findings are important for advancing real-time AR navigation systems that require both accuracy and efficiency in complex surgical environments. Potential applications include neurosurgery, liver surgery, and other procedures affected by soft tissue shifts. Limitations remain in terms of dataset diversity, limited anatomical coverage, and testing only on phantom models. Future work should extend evaluation to larger multi-center datasets, integrate real intraoperative imaging, and optimize computational performance for wider clinical translation.

